# Chemotherapy accelerated bone ageing is reversed by NMN

**DOI:** 10.1101/2025.10.13.679926

**Authors:** Maria B. Marinova, Romanthi Madawala, Wing-Hong Jonathan Ho, Vedran Lovric, Michael J. Bertoldo, Rema A. Oliver, Jayanthi Maniam, Margaret J. Morris, David A. Sinclair, Hayden A. Homer, Kirsty A. Walters, Jonathan H. Erlich, William R. Walsh, Robert B. Gilchrist, Lindsay E. Wu

## Abstract

Cancer patients face an array of long-term chronic diseases and accelerated biological ageing, due largely to the off-target effects of broadly cytotoxic chemotherapy drugs. This is especially a problem in children, where cancer survivors experience a subsequent high risk of bone mineral deficits and fractures, normally seen in the older population. Here, we model this to show that early-life treatment with a single dose of the commonly used chemotherapy cisplatin profoundly impairs late-life bone health, and that these bone deficits are completely resolved through treatment with the nicotinamide adenine dinucleotide (NAD^+^) precursor nicotinamide mononucleotide (NMN). While we had previously shown that this same strategy protects against chemotherapy induced female infertility, this maintenance of aged bone health appears to be unrelated to endocrine changes due to protection of the ovarian reserve. Rather, this is driven by altered phosphorus homeostasis and protection against renal damage, which otherwise increases parathyroid hormone secretion to mobilise calcium stores from bone. Overall, this work highlights a new approach for maintaining healthy bone ageing in cancer survivors.

## INTRODUCTION

Despite advances in oncology, cytotoxic chemotherapy drugs remain the mainstay of cancer treatment. These largely non-specific therapies have a high rate of adverse events with lifelong health consequences for those who survive cancer. In females, this can include depletion of the ovarian reserve, resulting in infertility and premature ovarian failure (POF) [1]. The loss of ovarian estrogen secretion has lifelong consequences [2] including metabolic dysfunction, neurocognitive disorders, cardiovascular disease and above all, an increased risk of osteoporosis. Strategies to maintain ovarian function in cancer patients could have profound impacts on the long-term health and quality of life of female cancer patients.

The diverse constellation of chronic disorders observed in cancer survivors include metabolic and cardiovascular disease, neurological disorders, bone disorders and cancers unrelated to the initial diagnosis [3, 4], together resembling a form of biological ageing, presenting a challenge to maintaining the long-term health of this patient cohort. Given this accelerated ageing phenotype, we recently investigated whether using strategies that had been applied to address ovarian ageing could also ameliorate ovarian damage caused by chemotherapy treatment [5]. We focused on the role of the enzyme cofactor nicotinamide adenine dinucleotide (NAD^+^), which we [6] and others [7-9] had previously shown to be a target for maintaining oocyte quality and fertility during reproductive ageing.

NAD^+^ plays another key role in slowing ovarian function by serving as a co-substrate of poly-ADP-ribose polymerase (PARP) enzymes in the DNA damage response [10], which is important to chemotherapy induced infertility [11]. These enzymes can be among the greatest consumers of NAD^+^ in the cell, depleting cellular NAD^+^ reserves to the point of impacting cell survival [12]. Further, many chemotherapy agents undergo metabolism by reductase enzymes that are fuelled by an NADPH cofactor, with NADP^+^ and NADPH levels dependent on the availability of NAD^+^ as a substrate for NAD^+^ kinase. Given this, we previously showed that boosting the NAD^+^ metabolome with its metabolic precursor nicotinamide mononucleotide (NMN) could ameliorate chemotherapy induced infertility in mice, with preservation of ovulation, follicle quality and breeding capacity [5]. Notably, this strategy did not compromise the oncological efficacy of chemotherapy against cancer, suggesting this could be a promising approach to maintain the fertility and long-term health of cancer survivors. In line with this, other groups also showed that NMN treatment could protect against the effects of chemotherapy treatment on oocyte quality [13, 14].

While fertility preservation strategies for cancer patients often focus on the cryopreservation of reproductive material for future fertility planning, another important aspect for oncofertility management is the future impacts on health from a loss in endocrine function [15]. In women, chemotherapy induced premature ovarian failure (POF) can lead to a loss in ovarian oestrogen production, which is essential for maintaining bone health. As a result, childhood cancer survivors can experience accelerated bone ageing, with an earlier risk of decreased bone mineral density, osteoporosis, fractures and frailty [16-20]. This is a major issue for cancer survivorship, particularly given that the bulk of the cohort of childhood cancer patients with improved survival are yet to reach old age.

Here, we sought to test whether early life chemotherapy treatment would impact accelerate bone ageing, and whether these changes to bone ageing would be reduced through the preservation of ovarian function through NAD^+^ repletion, as we had previously shown [5]. We found that a single dose of platinum chemotherapy in early life had profound impacts on late life bone ageing, which could be completely protected through treatment with the NAD^+^ precursor NMN. In contrast to our starting hypothesis, this was not mediated through a preservation of ovarian reserve and endocrine function, and was instead likely due to changes in renal toxicity, resulting in renal osteodystrophy.

## RESULTS

Previously, we showed that the NAD^+^ precursor NMN could prevent chemotherapy induced infertility in female mice that had been treated in early life with the platinum agent cisplatin (CDDP) [5], which is widely used in paediatric oncology for the treatment of solid tumours. This finding was likely related to slowing the rate of depletion of the ovarian reserve due to “follicular burnout”, a key mechanism in chemotherapy induced infertility [21]. In that study, we used a previously described model for chemotherapy induced infertility in the paediatric setting [5, 22, 23], whereby 7-day old mouse pups were treated with cisplatin (CDDP) alone. Two weeks later, at weaning, animals were allocated to drinking water supplemented with or without NMN, which was maintained throughout life to assess functional fertility, as measured by breeding trials (Fig. 1). As expected, CDDP treatment reduced the number of pups born per female, however this was rescued in animals that received subsequent NMN treatment, suggesting that even when delivered long after chemotherapy, NMN could maintain ovarian function.

**Figure 1.**
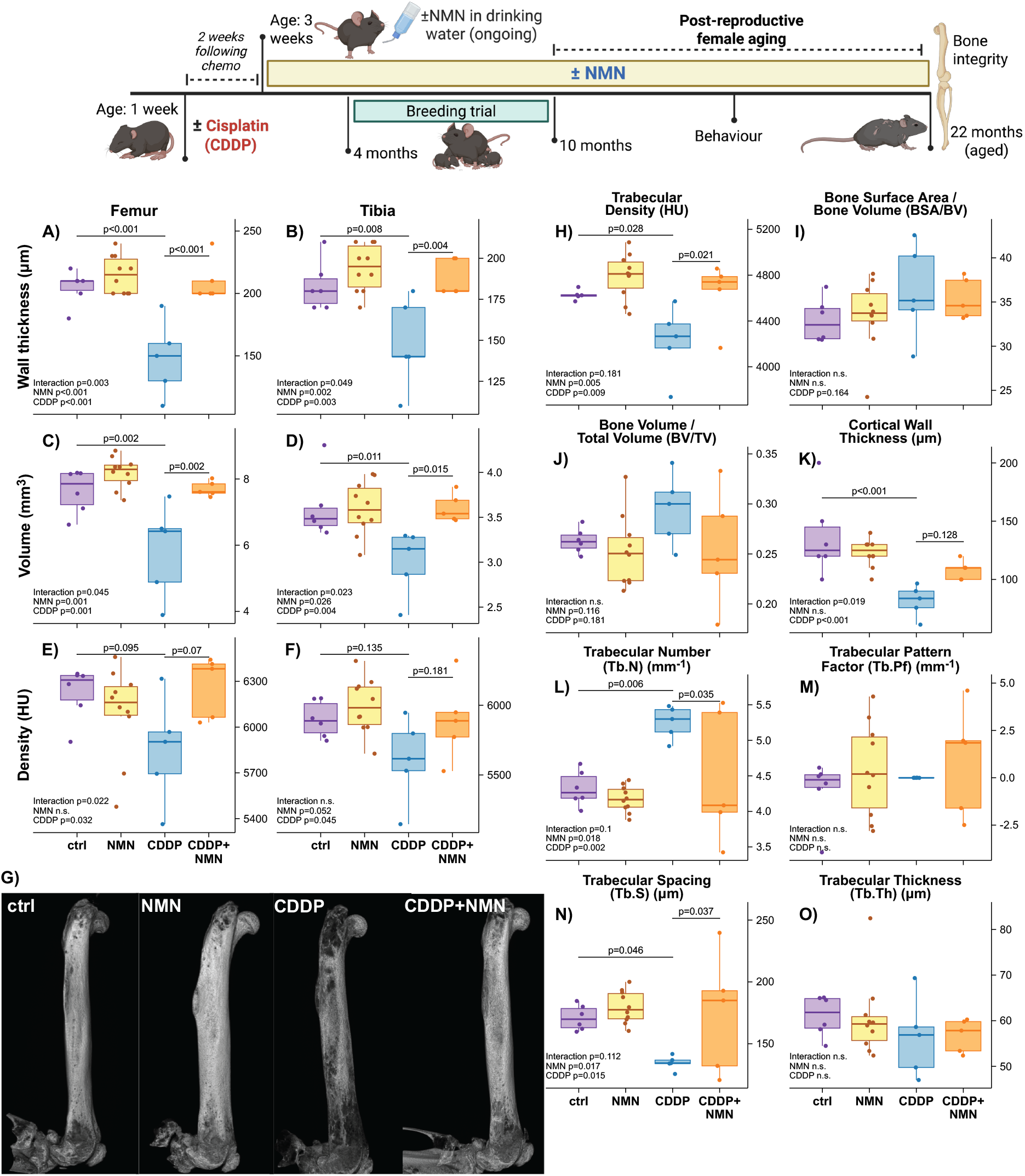
Late-life cortical bone structural changes following early-life chemotherapy treatment. Female mice treated in early life with CDDP (2 mg/kg) or saline control. Two weeks later, animals were maintained with or without NMN in drinking water (2 g/L) until euthanased for tissue collection at 22 months of age. Hindlimb femur diaphysis (A-C) and tibia cortical bone (D-F) were subject to µ-CT imaging from the midpoint of each bone for A, D) wall thickness, B, F) volume and C, F) bone density (Hounsfield units), with G) representative CT images of femurs shown. Femur trabecular bone (H-O) was also subject to µ-CT for H) bone density (Hounsfield units, HU), I) bone surface area to bone volume ratio, J) bone volume to total volume ratio, K) cortical wall thickness and trabecular parameters including L) trabecular number (Tb.N), M) pattern factor (Tb.Pf), N) spacing (Tb.S) and O) thickness (Tb.th). N=5-10/group as indicated by raw data points, p-values between groups are from Bonferroni-adjusted t-tests derived from estimated marginal means of linear model of NMN and CDDP treatment, with results of linear models indicated on each panel.

Aside from its implications for fertility, ovarian failure has profound endocrine impacts that impact systemic health. One key feature of this endocrine failure is impaired bone health, primarily due to the role of estrogen (E2) in regulating osteoblast and osteoclast function. Having observed protection against cisplatin-induced infertility, we proposed that this intervention would likely have profound impacts on late-life health and overall ageing, and in particular, bone function. To test this, following early-life cisplatin and lifelong NMN supplementation, animals were maintained until 22 months of age, close to the median lifespan of this mouse strain, to measure late-life bone structure and function (Fig. 1).

Hindleg femurs and tibias from this cohort of aged animals were subject to structural analyses of µ-CT imaging of femur diaphysis (Fig. 1A-C) and tibia cortical (Fig. 1E-F) bone. These imaging studies (Fig. 1G) revealed a profound reduction in cortical wall thickness (Fig. 1A, D), bone volume (Fig. 1B, E) and density (Fig. 1C, F) from CDDP treatment. Strikingly, each of these parameters were completely preserved in animals that subsequently received NMN treatment following early-life CDDP. To further investigate the internal microarchitecture of bones in these aged animals, trabecular bone morphology was also assessed by µ-CT, though due to the age of these animals, only trabecular bone from the femur metaphysis could be analysed (Fig. 1H-O), with very little trabecular bone from the tibia remaining at this age. As with the earlier parameters, trabecular density (Fig. 1H), cortical wall thickness (Fig. 1K) and trabecular spacing (Fig. 1N) were drastically reduced by early-life CDDP treatment, but were again rescued by subsequent NMN treatment. Interestingly, there was also a paradoxical increase in trabecular number in the CDDP group (Fig. 1L), however this well-known characteristic of endochondral ossification during severe osteoarthritis [24], as with the other parameters, was also reduced by subsequent NMN treatment.

We next sought to test whether this would translate into differences in mechanical integrity, including strength and flexibility. To assess this, femur (Fig. 2A, B) and tibia (Fig. 2C, D) were subject to the three-point bending test, a measure of the mechanical force that is required to break bones. These measures are highly clinically relevant during ageing, as decreasing bone strength can lead to a greater risk of bone fracture, requiring prolonged recovery in an older population. In line with µ-CT imaging results, both femur and tibia showed reductions in bone strength due to early-life CDDP treatment (Fig. 2A, C), though this was again normalised by subsequent NMN treatment in tibia (Fig. 2C). There was a trend towards reduced stiffness caused by NMN in the tibia, with a trend towards a reversal of this change by NMN treatment (Fig. 2D).

**Figure 2.**
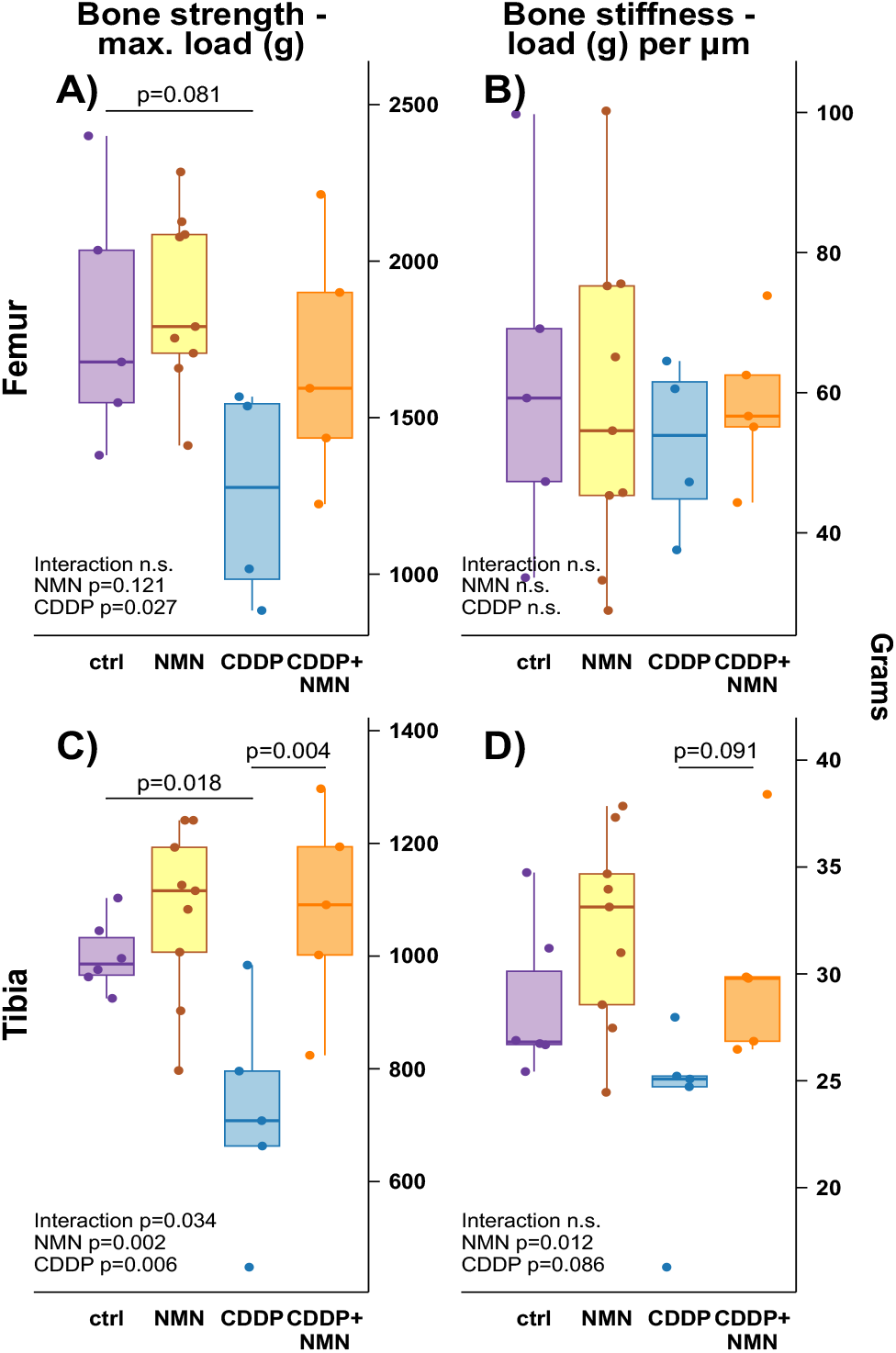
NMN maintains late life bone strength following early-life chemotherapy. Female mice treated in early life with CDDP and/or NMN, and bones collected from aged mice as per Fig 1. Bones were analysed by mechanical testing in a three-point bending setup to test A) femur strength; (B) femur stiffness; (C) tibia strength, and (D) tibia stiffness. N=4-9 per group as indicated by data points, p-values are from Bonferroni-adjusted t-tests derived from estimated marginal means of linear model of NMN and CDDP treatment, with results of linear models indicated on each panel.

Following µ-CT imaging (Fig. 1) and mechanical testing (Fig. 2), we next sought to investigate the microarchitecture of these aged bones through histology (Fig. 3). Cross-sections from femur metaphysis and diaphysis bone were stained and subject to blinded scoring for bone matrix organisation (Fig. 3A, B) and cortical porosity (Fig. 3C, D). Strikingly, cisplatin treatment led to profound changes in metaphysis bone, with severe matrix disorganisation and porosity (Fig. 3A, C). As with other parameters, these changes were rescued by NMN treatment. One unexpected observation from this histology was the pattern and severity of matrix disorganisation, porosity and chondrocyte infiltration caused by CDDP treatment (Fig. 3E). These changes do not resemble the changes to bone architecture that are classically observed during standard mouse models of osteoporosis, for example in the ovariectomised mouse model. Instead, they could resemble endochondral ossification, or alternatively, brown tumours from renal osteodystrophy.

**Figure 3.**
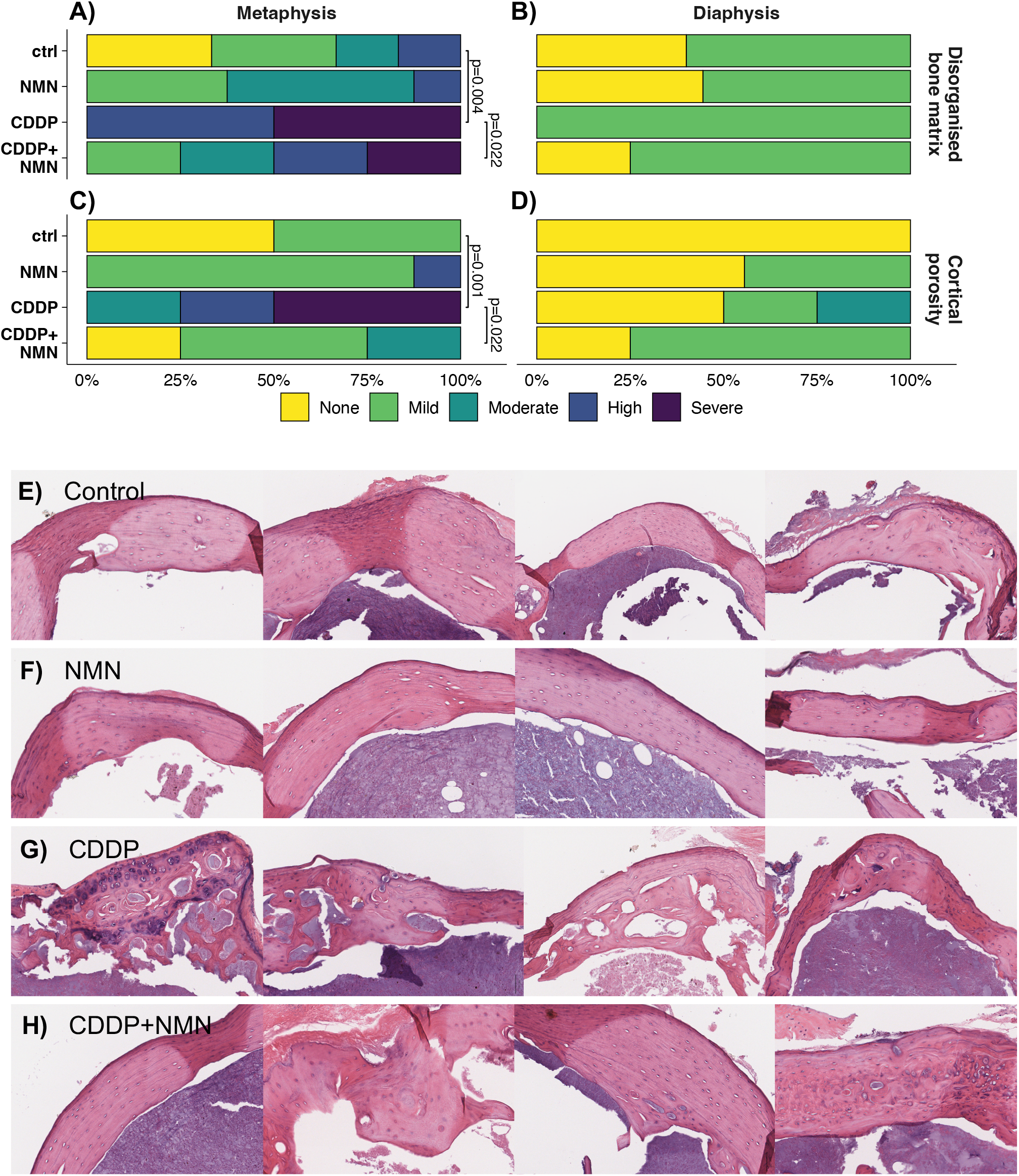
Bone matrix disorganisation following early-life cisplatin (CDDP) is rescued by NMN treatment. Female mice treated in early life with CDDP and/or NMN, and bones collected from aged mice as per Fig 1. Femurs were subject to histological assessment of A, B) bone matrix disorganisation and C, D) cortical porosity in A), C) metaphysis and B, D) diaphysis. E-H) Representative images of femur cross-sections at the metaphysis for E) control, F) NMN, G) CDDP and H) CDDP and NMN treated animals. Scale bar = 50 µm. N=4-10/group. Data analysed by ordinal logistic regression, p-values are from Bonferroni-adjusted t-tests derived from estimated marginal means of ordinal logistic regression model for CDDP and NMN treatment.

Together, these data present strong evidence of the ability of NMN to maintain bone integrity following chemotherapy treatment. Given the protection of bone by NMN treatment (Figs 1-3) and previous work on NMN in age-related [6, 7, 25-27] and chemotherapy-induced infertility [5, 13, 14], one obvious explanation for these findings is that bone loss following chemotherapy is due to declining estrogen secretion. To test this, we repeated cisplatin and NMN treatment and used mass spectrometry to assess a panel of sex steroids in a cohort of younger animals (Fig. 4), including testosterone, androstenedione, progesterone, estradiol (E2), estrone (E1) and follicle stimulating hormone (FSH) (Fig. 4A-F). In contrast to our hypothesis, these data did not support the idea that cisplatin caused endocrine insufficiency, with no change in any of these markers, including estradiol (Fig. 4D).

**Figure 4.**
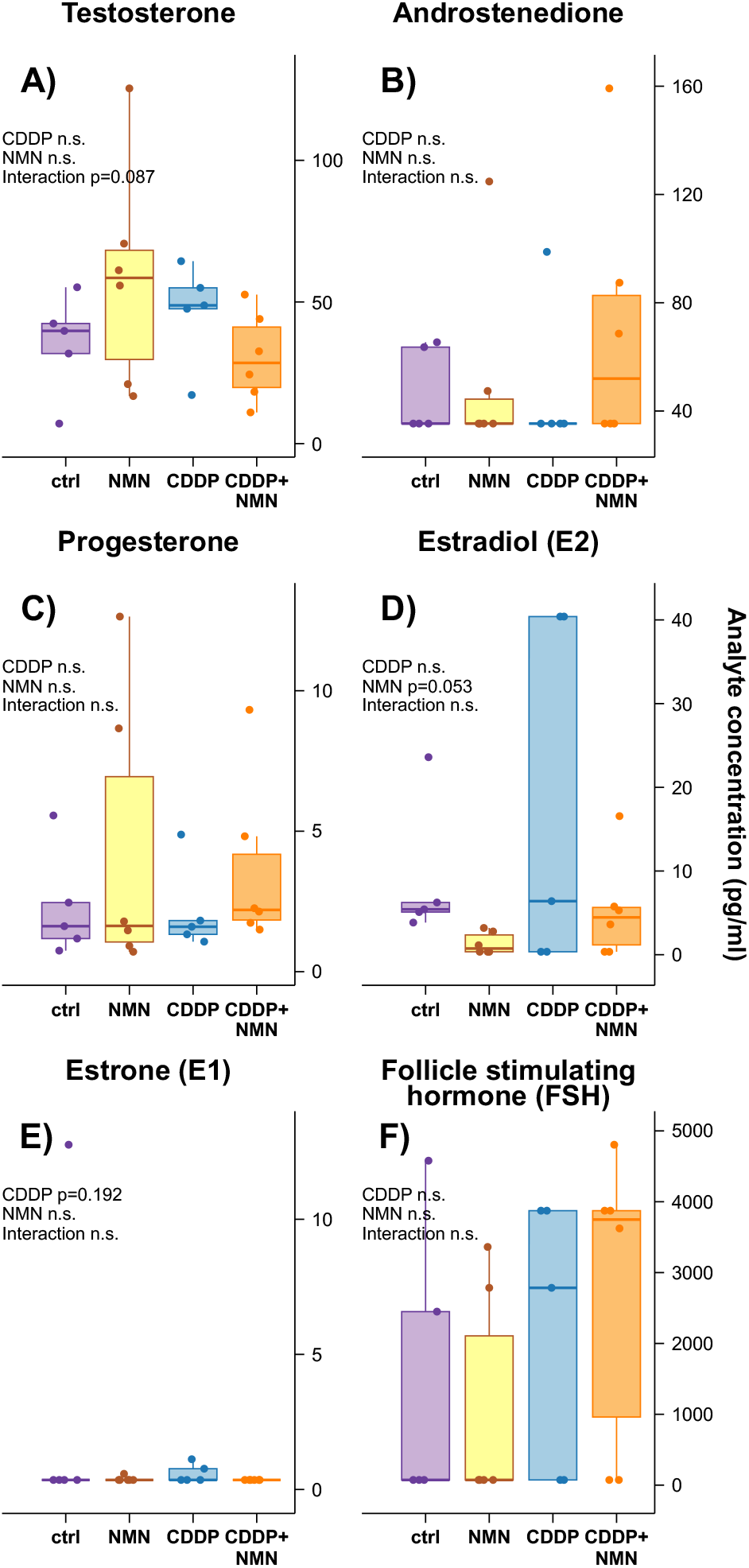
Sex hormone profile following early-life cisplatin. Mice were treated with CDDP and NMN as per Fig. 1, and serum collected at 10 weeks of age for measurements of A) testosterone, B) androstenedione, C) progesterone, D) estradiol (E2), E) estrone (E1) and F) follicle stimulating hormone (FSH). N=6 per group, data analysed by linear model of NMN and CDDP treatment, p-values from each factor and interaction as indicated.

These serum samples were collected from animals that were estrous cycle stage-matched, and so the lack of change is unlikely to be due to cyclic heterogeneity. While a strong indicator, these hormone levels do not necessarily reflect granular changes in the ovarian reserve, and could instead reflect changes in follicle function. Further, if ovarian changes and endocrine failure were to be the cause of altered late-life health, one would expect changes from an early age, given the stark degree of follicle atresia and burnout that occurs following a single dose of cisplatin. To investigate this, we measured the ovarian reserve in animals that were euthanased at three, six and ten weeks of age (Fig. 5, Extended Data Fig. 1). As expected, CDDP treatment caused early losses in primordial follicle reserve (Fig. 5A), transitory follicles (Fig. 5B) and primary follicles (Fig. 5C), though consistent with the “follicular burnout” hypothesis, increased the number of antral follicles in young, three-week old mice (Fig. 5D). Reductions in the primordial reserve persisted at later timepoints (Fig. 5A), however at no timepoint did we see subsequent NMN treatment ameliorating these reductions in the follicular reserve. We also collected bones from these younger animals for structural assessment of trabecular bone by µ-CT (Fig. 5F-J) – although we observed some early changes in trabecular pattern factor (Fig. 5H) and spacing (Fig. 5I) at three weeks of age, at six and ten weeks post-chemotherapy we did not observe the severe derangements in bone structure that were observed in aged mice (Fig. 1-3).

**Figure 5.**
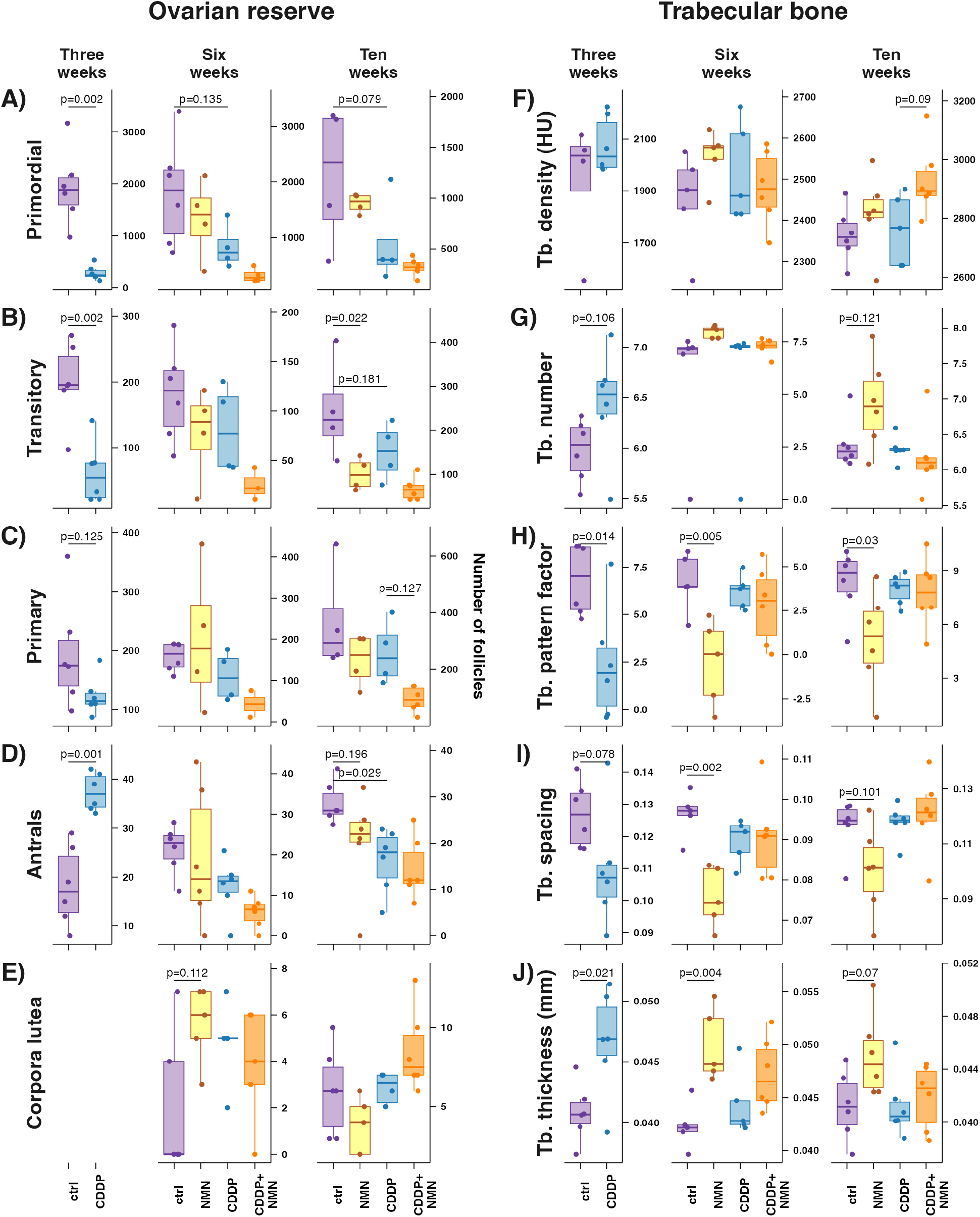
Time course of ovarian loss and bone microstructural changes. To test for a potential role of protection by NMN against early-life ovarian depletion from cisplatin (CDDP) as a cause of improved late-life bone health, tissues were collected following CDDP treatment as per Fig. 1. Tissues were collected at the start of NMN treatment at 3 weeks of age, with follow-on tissue collections at six and ten weeks of age. A-E) Ovaries were subject to stereology to assess reserves of A) primordial, B) transitory, C) primary and D) antral follicles, as well as E) corpora lutea remnants of ovulation. F-J) Femurs were collected for µ-CT analysis of trabecular bone microarchitecture, assessing F) density (Hounsfield units, HU), G) trabecular number, H) pattern factor, I) trabecular spacing and J) thickness. N=6 per group per, p-values are from Bonferroni-adjusted t-tests derived from estimated marginal means of linear model of NMN and CDDP treatment.

Together, the lack of changes in endocrine profile (Fig. 4), ovarian reserve (Fig. 5A-D) or early changes in bone structure (Fig. 5F-J), indicate that chemotherapy-induced ovarian follicle loss and endocrine failure are unlikely to be the cause of the severe phenotype that was observed in ageing (Fig. 1-3), suggesting other mechanisms may be involved in this bone phenotype. One clue to a potential mechanism for this phenotype is the striking changes in bone morphology observed with CDDP treatment (Fig. 3E), which was unlike that observed in osteoporosis induced by surgical removal of the ovaries. The striking morphology of cisplatin treated bones (Fig. 3) could also resemble renal osteodystrophy or osteitis fibrosa cystica, which occurs with kidney damage, also known as renal osteodystrophy, and is characterised by “brown tumours” that are reminiscent of the structures observed in cisplatin treated bone. In this condition, renal injury can lead to impaired phosphate excretion and impaired conversion of 25(OH)D into 1,25(OH)_2_D (vitamin D3; calcitriol), which is required for dietary calcium absorption. Changes in serum calcium and phosphate levels are detected by the parathyroid, which releases parathyroid hormone (PTH) and attempts to normalise serum Ca^2+^ levels through mobilising Ca^2+^ stores from bone, resulting in bone loss. Serum phosphate can also be detected by osteocytes, which release FGF23 to regulate phosphate excretion and further potentiate PTH secretion.

Platinum chemotherapy drugs such as cisplatin are well-known to induce renal toxicity, with cisplatin treatment acting as a common animal model for kidney damage. NAD^+^ precursors have previously been shown to offer protection against this damage [28-30], and one possibility is that this aged bone phenotype here is secondary to a rescue of renal toxicity. To test this, we first measured blood urea nitrogen (BUN), which is elevated in chronic kidney injury (Fig. 6A). There was an overall impact of cisplatin treatment on BUN, though the trend towards reduced BUN with cisplatin and NMN co-treatment was not significant. We also measured cystatin C, a serum marker of acute kidney injury (Fig. 6B), though no change was observed due to cisplatin treatment, likely due to the prolonged time between treatment and sample collection – despite this, there was a trend (p=0.111) towards reduced cystatin C with NMN. Renal injury is often associated with impaired phosphate excretion, which can lead to phosphate accumulation. We therefore measured serum inorganic phosphate (PO_4_) levels (Fig. 6C) which were unaffected by NMN or cisplatin treatment. While inorganic phosphate was unchanged (Fig. 6C), we also conducted atomic analysis of total, elemental phosphorus, which includes phosphorus atoms present in any species including inorganic phosphate (PO_4_), phospholipids, nucleic acids and others using inductively coupled plasma-mass spectrometry (ICP-MS). This revealed surprising changes in elemental phosphorus levels (Fig. 6D), which were almost halved by NMN treatment, regardless of cisplatin.

**Figure 6.**
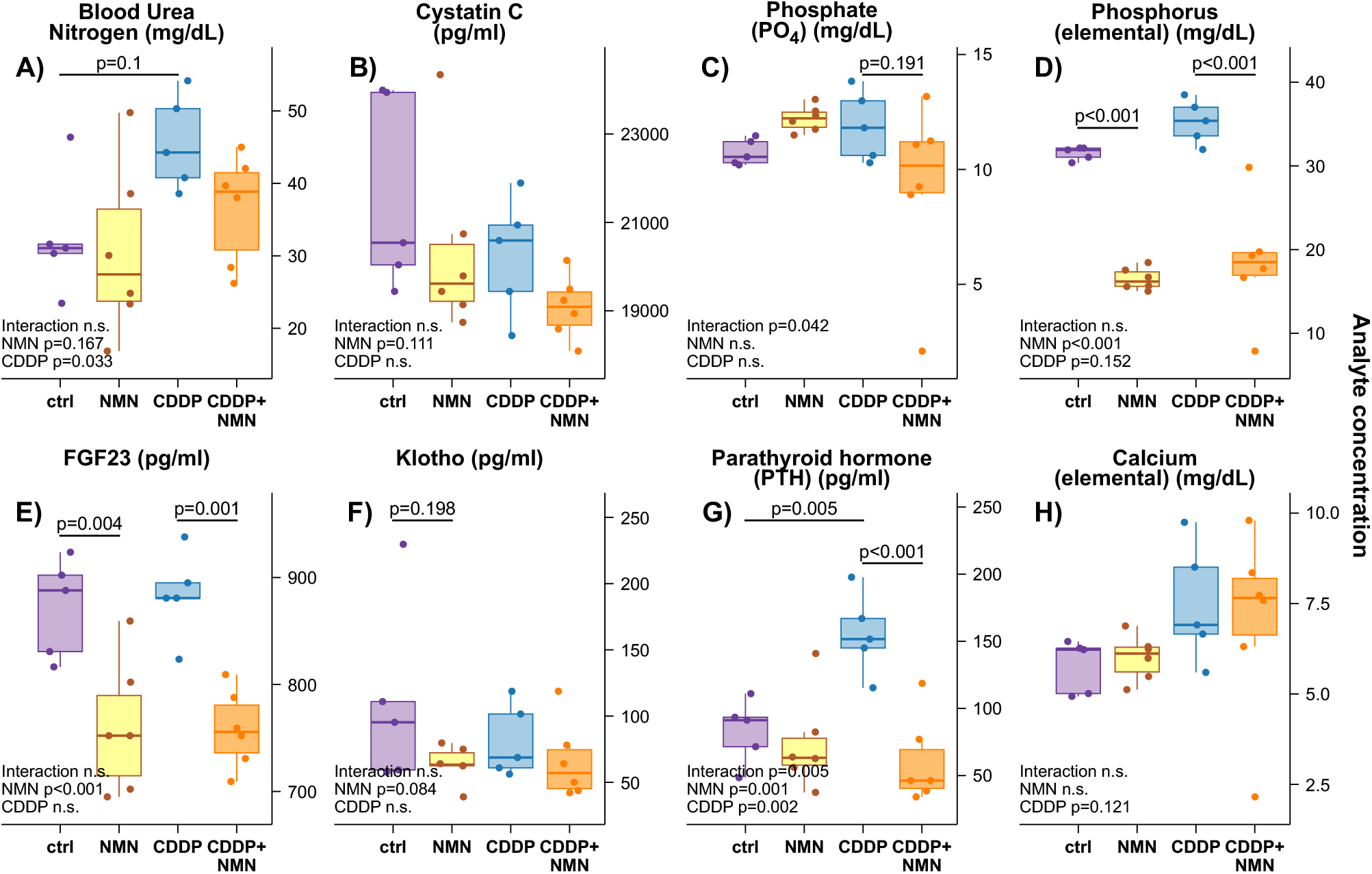
Renal damage, altered phosphate reabsorption and hyperparathyroidism as a cause of renal osteodystrophy. Mice were treated with CDDP and NMN as per Fig. 1, and serum collected at 10 weeks of age for measurements of the renal function markers A) blood urea nitrogen (BUN) and B) cystatin C. Renal damage that can impact the excretion of C) phosphate (PO_4_), which was not altered by these treatments, in contrast to D) elemental phosphorus, which encompasses both soluble, free phosphate, insoluble calcium phosphate precipitates, and organic phosphorus, such as phospholipids. Phosphate homeostasis can be regulated by the bone-derived hormone E) FGF23 and its coreceptor F) Klotho. These also regulate the production of G) parathyroid hormone (PTH), which maintain H) serum calcium levels and promote bone loss through the mobilisation of calcium from bone mineral stores. N=5-6 per group as indicated by data points, p-values are from Bonferroni-adjusted t-tests derived from estimated marginal means of linear model of NMN and CDDP treatment, with results of linear models indicated on each panel

Nicotinamide and NAD^+^ can impair phosphate reabsorption in renal tubules, resulting in increased phosphate excretion [31-34]. Given the need to maintain phosphate homeostasis within a tight range, the reduction in elemental phosphorus (Fig. 6D) likely reflects compensatory depletion of other pools of phosphorus, for example from organic phospholipids, to maintain serum levels of inorganic phosphate (Fig. 6C). Phosphate homeostasis is also regulated by osteocytes on the bone surface, which release FGF23 to promote renal phosphate excretion. In line with this, FGF23 was also reduced by NMN treatment (Fig. 6E), with a trend (p=0.084) towards a similar reduction in soluble forms of its coreceptor Klotho (Fig. 6F). Interestingly, recent work suggests that FGF23 secretion is regulated by organic phosphate in the form of glycerol-3-phosphate, rather than inorganic phosphate [35] – this reduction in FGF23 with NMN treatment (Fig. 6E) again supports the idea that the discrepancy between elemental phosphorus (Fig. 6D) and inorganic phosphate (Fig. 6C) is due to difference in the organic phosphate pool.

An important feature of renal injury is impaired Ca^2+^ homeostasis, due in part to the impaired excretion of phosphate, which can form insoluble calcium phosphate precipitates, and the reduced production of calcitriol, which regulates intestinal Ca^2+^ absorption. This impacts bone homeostasis through the production of parathyroid hormone (PTH), which acts to maintain Ca^2+^ levels in serum through mobilising reserves of Ca^2+^ from bone, and when chronically elevated in hyperparathyroidism, drives bone loss. While PTH production was previously thought to be driven by low Ca^2+^ levels, serum phosphate has emerged as a strong independent regulator of PTH, due to the inhibition of the calcium sensing receptor (CaSR) by physiologically relevant concentrations of serum phosphate [36]. Interestingly, serum PTH levels were more than doubled in the CDDP treated group, and this elevation was normalised to baseline by subsequent NMN treatment (Fig. 6G). This change in PTH secretion was likely due to altered phosphate homeostasis rather than changes in serum Ca^2+^, which was unchanged (Fig. 6H). Other circulating cytokines which were measured but did not change include growth hormone, prolactin, thyroid stimulating hormone and lutenising hormone (Extended Data Fig. 2). Together, these data support a model whereby a single dose of cisplatin early in life leads to chronic kidney disease, which impairs phosphate excretion, resulting in elevated PTH secretion (Fig. 6G) which mobilises bone stores of mineral calcium. This secondary hyperparathyroidism (Fig. 6G) can effectively maintain serum phosphate and calcium levels at the cost of impaired bone health (Fig. 1-3), resulting in late-life osteitis fibrosa cystica – a bone condition caused by chronic kidney disease and secondary hyperparathyroidism. With NMN treatment, impaired phosphate reuptake results in ongoing urinary phosphate excretion, preventing the increase in PTH levels that mobilise Ca^2+^ from bone mineral stores, thereby preserving bone integrity.

Given our original hypothesis around ovarian depletion driving endocrine failure and impaired late-life health, we also assessed whether this would impact aspects of behaviour, which would link to the psychological changes that occur with menopause in humans. Animals were subject to the elevated plus maze, which is intended to measure anxiety-like behaviour (Fig. 7). To our surprise, animals treated with CDDP alone spent more time in the open arm (Fig. 7A) – an effect that was again ameliorated by NMN co-administration.

**Figure 7.**
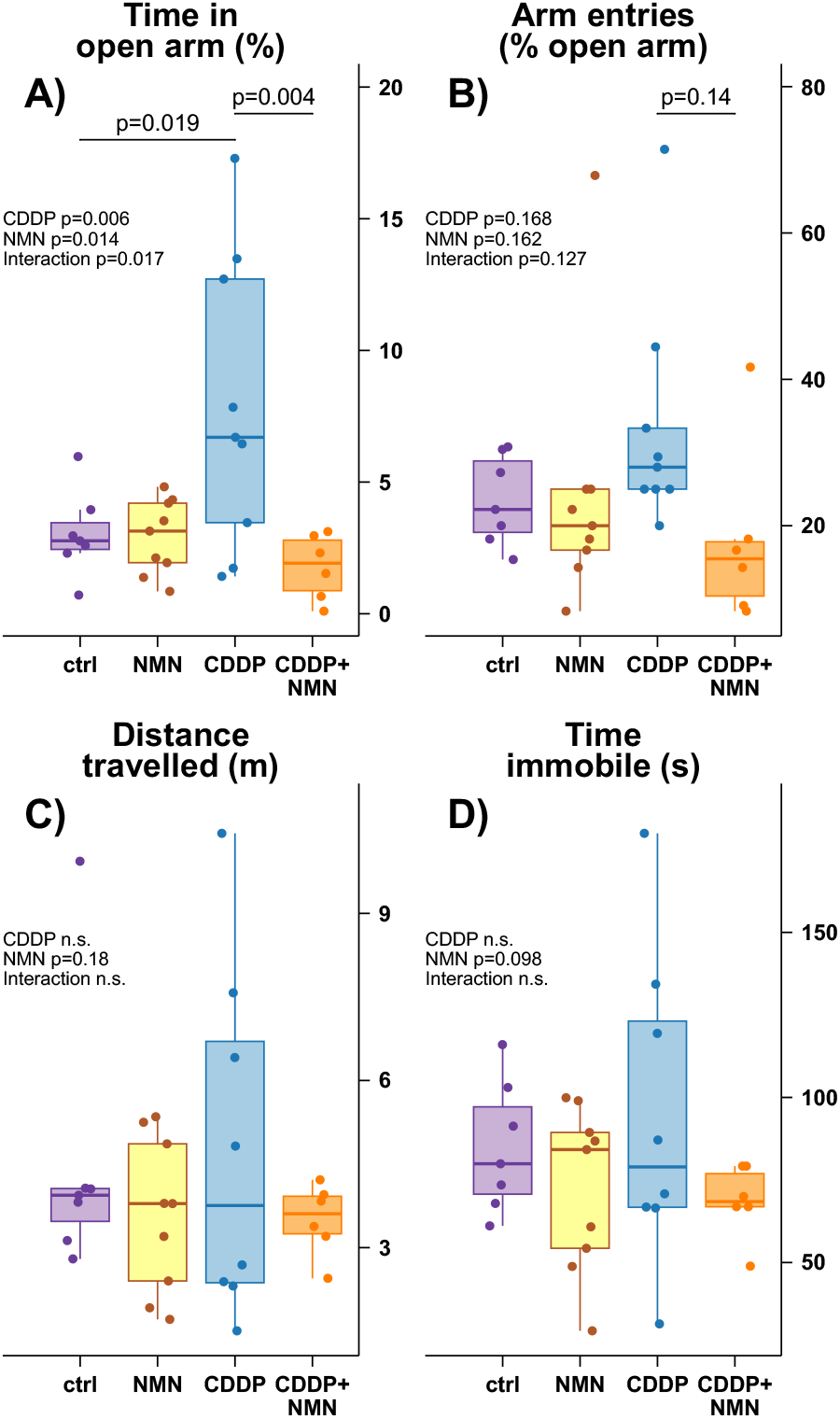
Behavioural impacts of cisplatin and NMN. At 13 months of age, animals treated as per Fig. 1 were subject to the open field test to test for potential impacts on anxiety-like behaviour. This measured A) the time spent in the open arm, B) the percentage of arm entries into the open arm, C) overall distance travelled, and D) time spent immobile. The increased proportion of time spent in the open arm with cisplatin treatment is evidence of reduced anxiety-like behaviour, which is normalised to baseline levels by subsequent NMN treatment. N=6-9 per group as indicated by data points, p-values are from Bonferroni-adjusted t-tests derived from estimated marginal means of linear model of NMN and CDDP treatment, with results of linear models indicated on each panel.

## DISCUSSION

Here, we present a new strategy to prevent bone loss in cancer patients following chemotherapy treatment, in the form of treatment with the NAD^+^ precursor NMN – an intervention that prevents chemotherapy induced in mice infertility, even when delivered weeks after chemotherapy [5]. This treatment shows promise as a strategy for use in the context of cancer, as we previously showed that NMN does not reduce chemotherapeutic efficacy against tumour growth [5]. This study emphasises the importance of the biology of ageing in maintaining long-term health in cancer survivors, with other work demonstrating a role for senescence in chemotherapy induced bone loss [37].

This finding has implications for the health of paediatric cancer survivors, who face long-term health consequences resulting from cytotoxic chemotherapy treatment, with a high rate of chronic health conditions that resemble accelerated biological ageing [4]. In female patients, these treatments can result in infertility [2] due to premature depletion of the ovarian reserve [11]. Aside from the devastating social and health consequences of infertility and its impacts on family planning [1], these patients are also at a high risk of late-life adverse health effects, with osteoporosis [38], impaired bone mineral density and increased fracture risk representing key long-term health challenges [20, 39]. Notably, childhood cancer survivors experience a five-fold increase in musculoskeletal damage [40]. Similarly, breast cancer treatment results in impaired bone health, notably due to long-term maintenance therapy following treatment [41]. This risk of bone loss presents one of the greatest risks to late-life health in female cancer survivors, with the current treatment paradigm based on the assumption that bone loss is a secondary effect from premature ovarian failure and estrogen insufficiency. Despite this, there is no correlation between bone mineral density and E2 levels following chemotherapy [42]. This lack of correlation between bone mineral density and E2 in patients aligns with our findings, which in contrast to our initial hypothesis, suggest that bone loss following cisplatin treatment (Fig. 1-3) is unrelated to changes in sex steroid levels (Fig. 4) or the ovarian reserve (Fig. 5).

Rather than being due to a protection of ovarian function, this late-life bone phenotype (Fig. 1-3) was instead likely related to the well-characterised nephrotoxicity of cisplatin, which resulted in secondary hyperparathyroidism (Fig. 6H). PTH secretion regulates serum Ca_2+_ homeostasis, however this is achieved by mobilising bone mineral stores of Ca^2+^. Over time, the ongoing depletion of bone mineral to yield Ca^2+^ can impair bone health, and hyperparathyroidism is associated with the bone condition osteitis cystica fibrosa [43]. While many chemotherapy agents are associated with nephrotoxicity and chronic kidney disease [44, 45], whether this directly translates to secondary hyperparathyroidism or renal osteodystrophy is less clear. The only study to have measured the impact of cisplatin on PTH [45] levels used a small number of case studies, however this demonstrated that PTH levels were elevated two to three times over baseline levels following treatment. Future work should aim to investigate whether the elevated risk of impaired bone function in childhood cancer survivors [16-20] is related to chronic kidney damage and secondary changes in PTH levels.

Clinically, hyperparathyroidism and renal osteodystrophy can be resolved by surgical removal of the parathyroid (parathyroidectomy), or treatment with agents including the calcium mimetic cinacalcet to activate the calcium receptor CaSR, phosphate binders to reduce phosphate signalling, or vitamin D analogues to promote dietary Ca^2+^ absorption. Given the widespread interest in NAD^+^ precursors such as NMN for other conditions including in ageing and their ongoing clinical development, these compounds may be a convenient option for testing in a clinical setting for the treatment of hyperparathyroidism, which could be relevant to the later-life health of cancer survivors.

One important observation from this work was the striking reduction in serum phosphorus levels, which were almost halved by NMN treatment in both vehicle and cisplatin treated animals (Fig. 6D). NMN can undergo metabolism by hepatic, microbial and enzymatic pathways [46-50] to release free nicotinamide, which along with NAD^+^ can inhibit phosphate reabsorption in nephrons [31-34], resulting in the elevated urinary excretion of phosphate. Notably, this change in elemental phosphorus (Fig. 6D) was not correlated with levels of free inorganic phosphate (PO_4_) (Fig. 6C), suggesting that phosphate homeostasis is tightly maintained via buffering from some other unknown pool of elemental phosphorus – for example, from phospholipid species such as phosphatidylcholine. In support of the idea that phosphate homeostasis is tightly maintained, decreased elemental phosphorus levels with NMN treatment (Fig. 6D) were accompanied by decreased serum FGF23, a bone-derived hormone released in response to elevated phosphate levels which signals to promote renal phosphate excretion (Fig. 6E). Together, these findings support a scenario whereby NMN promotes phosphate excretion, resulting in compensatory reductions in FGF23 (Fig. 6E) and the mobilisation of serum pools of elemental phosphorus (Fig. 6D) present in species such as phospholipids, to maintain serum phosphate (Fig. 6C) homeostasis. An alternative explanation for the disconnect between inorganic phosphate (Fig. 6C) and elemental phosphorus (Fig. 6D) could be that if NMN promotes the urinary excretion of phosphate, there would be decreased formation of insoluble calcium phosphate precipitates. These colloidal precipitates would be detected by elemental analysis, whereby samples undergo complete deposition in concentrated nitric acid prior to analysis, but masked from the malachite green assay for soluble inorganic phosphate that was used here. If so, one would expect a similar change in elemental calcium levels, due to the fixed stoichiometry of calcium to phosphate in calcium hydroxyapatite or amorphous insoluble calcium phosphate, however elemental Ca_2+_ levels (Fig. 6G) were not changed in line with elemental phosphorus (Fig. 6D). The possibility that NMN impacts the formation of insoluble calcium phosphate precipitates through its promotion of phosphate excretion is not inconsistent with these results, as differences in the formation of insoluble precipitates could be masked by their deposition on vascular walls, leading to vascular calcification. It still would not, however, explain the disconnect between elemental phosphorus (Fig. 6D) and inorganic phosphate in serum (Fig. 6C), and future work should aim to characterise the identity of this “missing pool” of elemental phosphorus.

The direct impairment of renal phosphate reuptake by NMN or its metabolites, resulting in urinary phosphate excretion, is a likely explanation for our result, however the drastic reductions in serum levels of elemental phosphorus have, to our knowledge, not been described with NMN treatment. Given that long-term treatment with NMN and NR is being investigated as a strategy to maintain late-life health, it will be important to establish whether phosphaturia occurs during chronic treatment with these NAD^+^ precursors, and whether this can be leveraged as a strategy to replace existing phosphate binding drugs, or for the treatment of hyperparathyroidism. Overall, it appears that reductions in the elemental phosphorus pool with NMN treatment are unrelated to cisplatin injury, however the net effect of this treatment overlaid onto injury induced by cisplatin is to overcome hyperparathyroidism (Fig. 6H) and completely ameliorate renal osteodystrophy, with striking protection against deranged bone morphology (Fig. 3), weakness (Fig. 2) and structural changes (Fig. 1).

If true, these findings could provide a mechanism that unites two disparate models in the biology of ageing. One of the earliest genes thought to play a role in biological ageing encodes for the transmembrane protein Klotho, which is a co-receptor for FGF23. Mutation in Klotho results in shortened lifespan and a progeria phenotype [51], while its transgenic overexpression extend lifespan [52]. Klotho knockout mice have elevated serum phosphate and extremely high circulating FGF23 levels, however deletion of the renal phosphate transporter NaPi2a in Klotho knockout mice rescues hyperphosphatemia and results in normal development [53]. This rescue of the progeria phenotype in Klotho knockouts by NaPi2a deletion can then be abrogated by feeding these animals a high phosphate diet [54]. Together, these results have led to a “phosphate toxicity” model of ageing [54, 55] that is mediated by the Klotho-FGF23 axis. Independently of this, there has been strong interest in an “NAD^+^-centric” model of ageing, whereby declining levels of the enzyme cofactor NAD^+^ during ageing are thought to impair redox cofactor-dependent cellular metabolism and the activity of epigenetic and DNA repair enzymes such as the sirtuins and poly-ADP-ribose polymerases (PARPs). This has led to the widespread investigation and development of NAD^+^ biosynthetic precursors such as NMN (as used here) and nicotinamide riboside (NR) in an attempt to extend lifespan and for the treatment of age-related disorders [56-59], infertility [5-7, 26, 27] and inflammation, resulting in clinical trials of these compounds. Given the well-established role of the NAD^+^ metabolite nicotinamide in promoting phosphate excretion [34, 60] and preventing its intestinal absorption [33, 61], it may be possible that some of the beneficial effects of treatment with NAD^+^ precursors could relate to a lowering of the phosphate burden, which is otherwise regulated by FGF23 and Klotho. This link will likely have been missed in previous work due to the use of assays that measure soluble, inorganic phosphate using the malachite green assay, rather than the elemental analysis used here (Fig. 6D).

In this proposed model of crossover between the Klotho-FGF23 driven model of phosphate toxicity and the NAD^+^ deficiency model of ageing, treatment with NAD^+^ precursors could also be expected to improve cardiovascular function through reducing the incidence of tissue calcification. Increased phosphate excretion through inhibition of its renal reuptake by NAD^+^ metabolites would be expected to reduce the formation of insoluble calcium phosphate precipitates, which can be deposited on vascular walls and leading to vascular calcification and stiffness. Under this scenario, previous findings around the role of NAD^+^ homeostasis in maintaining vascular health during ageing [62] could also be explained by differences in vascular calcification. This model would also be relevant to the proposed role of vascular stiffness in brain ageing and the vascular dementia model [63, 64]. Future investigations should aim to test the impact of NMN treatment on the rate of formation of calcium phosphate precipitates, and vascular calcification in ageing.

This study also revealed changes in animal behaviour, with a potential reduction in anxiety-like behaviour in animals treated with CDDP alone (Fig. 7A, B). Previous work has shown a potential role for PTH signalling in anxiety, as genetic deletion of the PTH receptor PTHR2 and its peptide ligand TIP39 results in increased anxiety-like-like behaviour, while treatment with TIP39 reduces this [65-67]. Given this, the behavioural changes observed here may be a secondary consequence of increased PTH levels, rather than a direct, lasting effect of early-life cisplatin treatment.

Several additional experiments could be carried out to support our conclusions. Our interpretation that the striking protection against cisplatin induced bone-loss is independent of ovarian function is based on our finding that NMN treatment does not impact reductions in the follicle reserve due to cisplatin treatment (Fig. 5A), with a corresponding lack of change in estrogen (E2) levels (Fig. 4D). To verify whether the protection of bone function is truly independent of ovarian function – which is especially relevant given our previous finding that NMN can maintain fertility – future work should aim to replicate this study in ovariectomised mice, which will be estrogen deficient due to the surgical removal of ovaries. A similar experiment has been undertaken previously, showing that NMN treatment can protect against osteoporosis in ovariectomised mice [68], again supporting our conclusion that this protection is independent of endocrine changes, however it is unclear whether cisplatin treatment will result in additive effects to ovariectomy on bone health. If NMN can maintain protection of bone following cisplatin treatment in ovariectomised mice, this would be strong evidence for an alternate pathway, for example the renal osteodystrophy model that we propose here. Alternatively, a similar experiment could be performed using pharmacological treatment with aromatase inhibitors that block the production of estrogen. If these results show that this protection is independent of estrogen production, these findings could also be relevant to males, who would similarly be at risk of cisplatin induced nephrotoxicity, with potential impacts on bone health.

It would be of interest to determine whether NMN and cisplatin interact directly with bone, through in vitro studies of osteoblast function. To confirm our model that these changes in bone function are due to altered renal function and hyperparathyroidism, future experiments could use pharmacological antagonists of the parathyroid hormone receptor to test whether these too can prevent the profound loss in aged bone integrity that we see here. Finally, future work could aim to confirm changes in renal function using additional parameters such as glomerular filtration rate and phosphate reabsorption, though the impact of NAD^+^ homeostasis on cisplatin induced renal toxicity has been demonstrated elsewhere [69].

Together, this work is relevant to the long-term survival of cancer patients, suggesting a potential new strategy to maintain bone health, and highlighting pathologies that may be relevant to the healthy ageing of this population. This may be especially relevant to those receiving early-life chemotherapy treatment, and while the use of NAD^+^ precursors such as NMN will require ongoing clinical development, it will be interesting to see whether patients who received platinum-based chemotherapy and experienced lasting nephropathy will also be at risk of elevated PTH and impaired bone health.

**Extended Data Figure 1.**
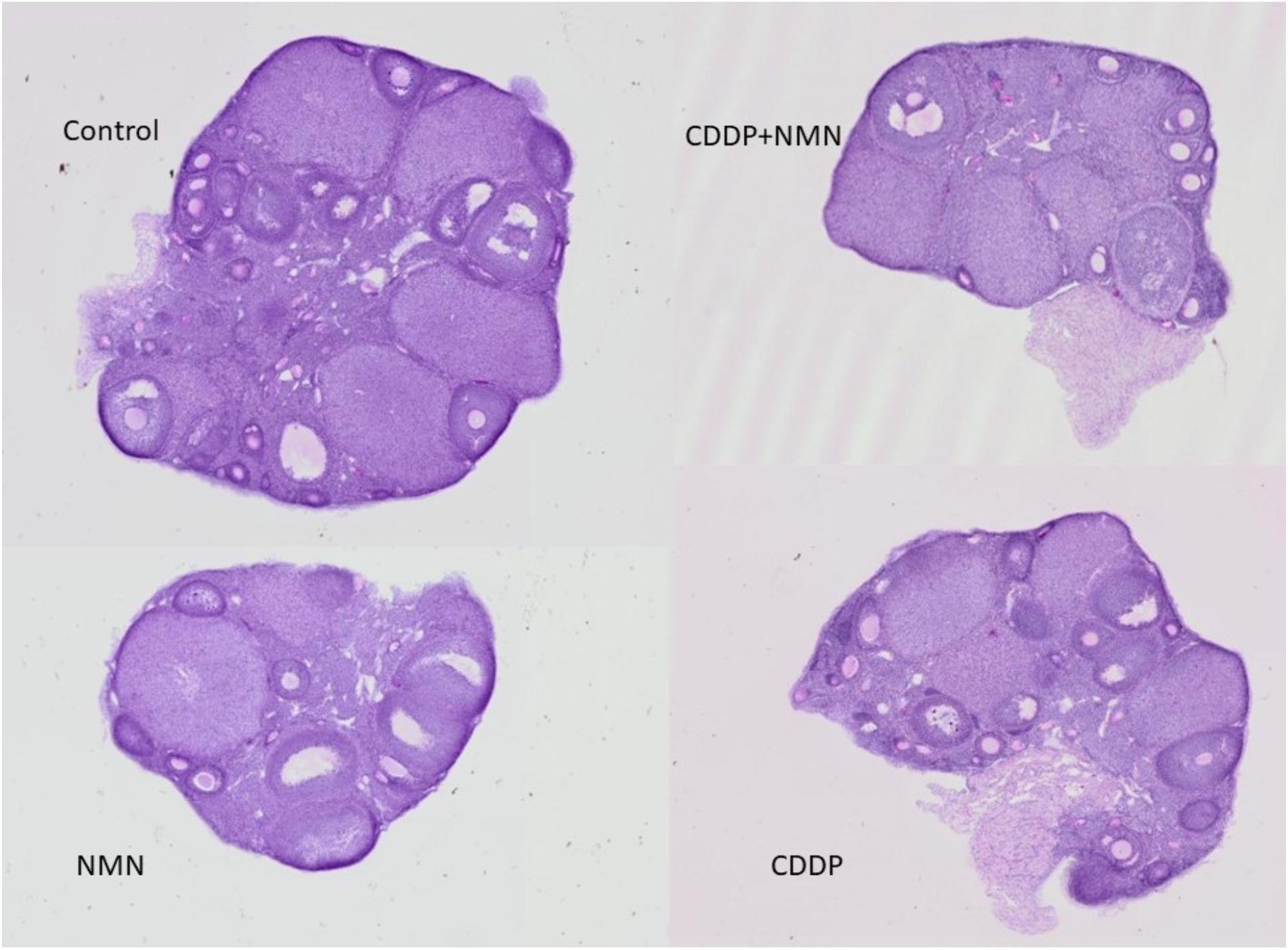
Representative ovarian histology. Ovaries were collected from ten-week-old female mice used for assessment of the ovarian reserve as shown in Fig. 5A-E, following early-life treatment with cisplatin and NMN as described in Fig. 1. Estrous cycling was tracked to ensure tissue collection of ovaries at diestrous, representative resin sections shown here.

**Extended Data Figure 2.**
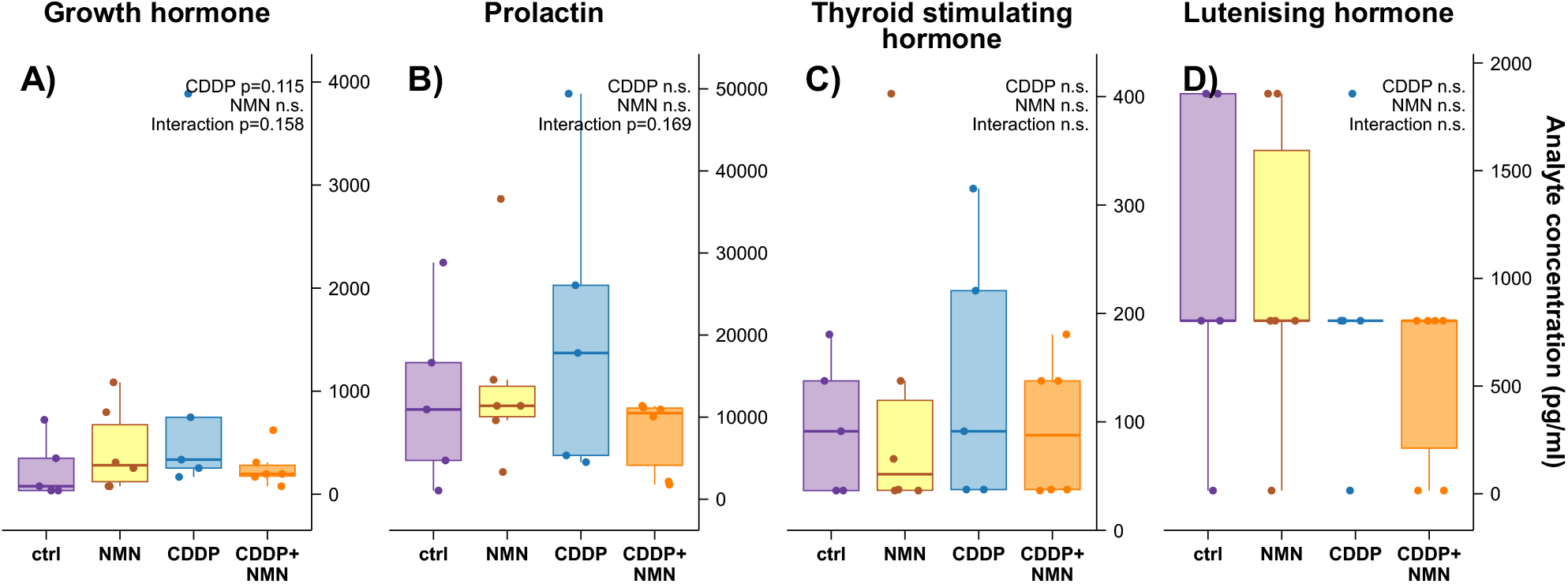
Additional serum biomarkers related to Figure 6. Serum collected from ten-week-old females as described in Figure 6 was analysed for A) growth hormone, B) prolactin, C) thyroid stimulating hormone and D) lutenising hormone.

## METHODS

### Animals

#### Ethics

Animal experiments were carried out with prior approval of the UNSW Animal Care and Ethics Committee, which operates under guidelines from the National Health and Medical Research Council of Australia.

#### Housing

Animal experiments were as described previously [5]. C57/BL6J^Aus/b^ mice were maintained in the UNSW Biological Resource Centre in individually ventilated cages at 22°C and 80% humidity, at a density of between two to five per cage, with *ad libitum* access to food and water. Drinking water was acidified with HCl to pH 3 to prevent microbial growth, this same water was used as a vehicle for dissolving NMN (2 g/L). Animals were maintained on standard chow diet obtained from Gordon’s Specialty Stock Feeds (Yanderra, NSW Australia) as described previously [70], briefly, this diet contained 8% calories from fat, 21% from protein, and 71% from carbohydrates, with an energy density of 2.6 kcal/g. The UNSW animal house maintained a 12 hr light/dark cycle, with lights on at 0700 and off at 1900.

Bone samples from aged animals subject to early life cisplatin and/or NMN treatment were taken from female mice previously subject to the breeding trial described in our previous work [5]. In this study, seven-day old pups were injected with cisplatin or saline control in the absence of NMN, and then weaned as per normal at 3 weeks of age onto standard acidified drinking water with or without the addition of NMN (2 g/L). At 16 weeks of age, animals were subject to a breeding trial. Following completion of the breeding trial described in that paper, animals were maintained until 22 months of age, then sacrificed to collect samples for bone analysis, as described below.

#### NMN and chemotherapy treatments

Mice were maintained on standard acidified drinking water in the presence or absence of NMN (Geneharbor Hong Kong Technologies Ltd, Hong Kong) at 2g/L, with water bottles changed twice weekly to prevent microbial growth, with treatment maintained until the end of the experiment. Cisplatin (Enzo, Switzerland) was administered at 2 mg/kg dissolved in saline – we were careful to avoid the use of DMSO, which can reduce the effectiveness of cisplatin by altering the coordination of Pt [71].

### Bone analyses

#### µ-CT bone imaging

Following sacrifice, whole hind right legs were preserved at –80°C. These samples were later thawed, cleaned of tissue and subjected to µ-computed tomography (CT) imaging, using a MILabs U-CT. Before and after the procedure they were kept in PBS and not allowed to dry out. Both the femur and the tibia were imaged. The images were taken using ultra focus magnification mode with a scan angle of 360°. The scan mode was set to accurate and the angle step degree was 0.25, with an exposure time of 75 ms. The X-ray voltage was 50 kV and current 0.21 mA. A double bed was used to image both the femur and tibia together. 3D reconstruction was performed using the PMOD software package (PMOD Technologies Ltd., Zurich, Switzerland) at 20-µm voxel size with a Hann and Gaussian volume filter (FWHM 24 micron). After DICOM conversion, analysis was performed using the Inveon software package.

#### Bone morphometry

Bone morphometry assessment of the µ-CT bone reconstructions was performed according to guidelines [72]. CT scans were analysed using 400 sections/bone of the femoral diaphysis for cortical bone −200 above and 200 below the middle section, 300 sections/bone of tibial diaphysis for cortical bone, and 100 sections/bone of femoral epiphysis and metaphysis for trabecular bone measurements. The selection of the material of interest was performed manually on the Inveon workspace software.

Every 5^th^ section for trabecular measurements and every 10^th^ section for cortical measurements were manually outlined in a blinded fashion, and the sections between those manually selected, were interpolated using 3D visualisation software (Inveon Research Workplace, Siemens, Germany), which were then reviewed and corrected. The selected areas were used to calculate bone morphology outcomes including; bone volume/total volume (BV/TV), bone surface area/bone volume (BSA/BV), trabecular number, thickness, spacing and pattern factor, as well as cortical and trabecular density and volume. In addition to these, cortical volume, density, and wall thickness were also measured from both the femur and tibia. Relative density was measured in Hounsfield units. Default settings were used for analysis, and calibrated using water (0 HU) and air (−1000 HU). These are all factory calibrated and not user calibration.

#### Mechanical testing: Three-point bending

Following µ-CT imaging, three-point mechanical testing of bones was performed as previously described [73]. Briefly, long bones were stripped of surrounding soft tissue and kept moist in PBS to prevent dehydration. Femurs were placed in a specifically designed holder to fit the length of the bone. The two holding points were set up 10 mm apart for the femur, while for the longer tibias, this length was adjusted to 11.2 mm. Three point-bending involves applying pressure to the bones to the point of mechanical failure, with the bone sitting on two support points as the third point, the lever, pushes on top of it (Extended Data Fig. 3). The force that is required to break each bone is recorded as well as the slope measured in force over time. The force required to break the bone is measured in grams of maximum load, while stiffness is measured by the slope of force over the time taken to break the bone completely from the point of initial contact. The slope provides a measure of bone flexibility prior to the break. The measured amount of force was deployed perpendicular to the middle of the anterior side of the diaphysis. The bones were loaded until the point of mechanical failure. The ramp amplitude was 1000 µm and ramp velocity 1 was 50 µm/sec, to generate a curve with a relaxation time of 1 sec. Ramp velocity 2 and fixed relaxation time 2 were 50 µm/sec and 4 sec as follows. Position and load vs time were monitored. Data output was analysed by the Mach-1 analysis software package (Biosyntech Inc, Netherlands). Not all samples were subject to mechanical testing, as µ-CT imaging revealed that some bones were broken prior to mechanical testing, likely due to

Prior to mechanical testing, µ-CT imaging revealed fractures on some samples, which hence could not be used for mechanical testing. The actual numbers used for analysis were as follows:

- Vehicle control: six femurs, one damaged near the head so not used for trabecular measurement in CT, and six tibias
- CDDP alone: five femurs, one damaged so not used for mechanical testing, and five tibias
- CDDP and NMN: five femurs, five tibias
- NMN, vehicle: ten femurs, one damaged so not used for mechanical testing, ten tibias, one damaged and not used for mechanical testing.

**Extended Data Figure 3.**
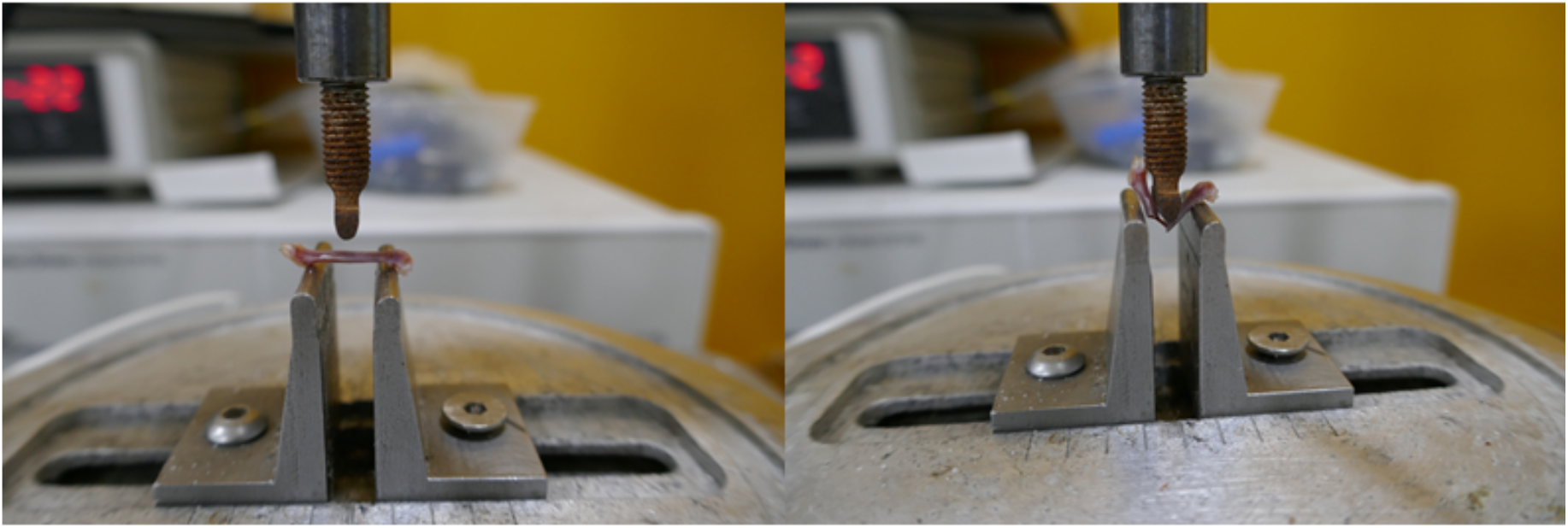
Mechanical three-point bending. Experimental setup for measuring the mechanical parameters of long bones in a three-point bending experiment. These measurements provided the mechanical strength and stiffness of bones shown in Fig. 2.

#### Histology

Following mechanical testing, bones were stored at –80°C. For histological analysis, samples were thawed at room temperature and placed in histology cassettes, then fixed in 10% neutral buffered formalin on a shaker at room temperature for 72 hrs. They were then decalcified in a 10% formic acid-formalin solution. The femurs were oriented to be cut axially and embedded in paraffin, with 5 µm sections cut and stained with Harris hematoxylin and eosin.

#### Histological assessment of bone structure

Matrix porosity and organization were assessed in a blinded fashion on a relative five-grade scale (0-4), where zero indicates a healthy bone structure and four signals the highest degree of damage. Blinding was achieved by sample de-identification: all labels indicative of treatment were substituted with a random number (sample ID). Sample IDs were only matched to animals from specific treatment groups upon completion of data analysis.

Matrix porosity was assessed under a 2.5-40x light microscope. Matrix organization was analysed under polarized light using a polarized light filter. Both assessments were scored under the following descriptions:

- 0 - no abnormal phenotype (healthy)
- 1 - very mild matrix porosity/matrix disorganisation phenotype
- 2 - medium matrix porosity/ matrix disorganisation phenotype
- 3 - high matrix porosity/ matrix disorganisation phenotype
- 4 - severe matrix porosity/ matrix disorganisation phenotype

Representative examples of each of these categories are shown in Extended Data Figure 4.

**Extended Data Figure 4.**
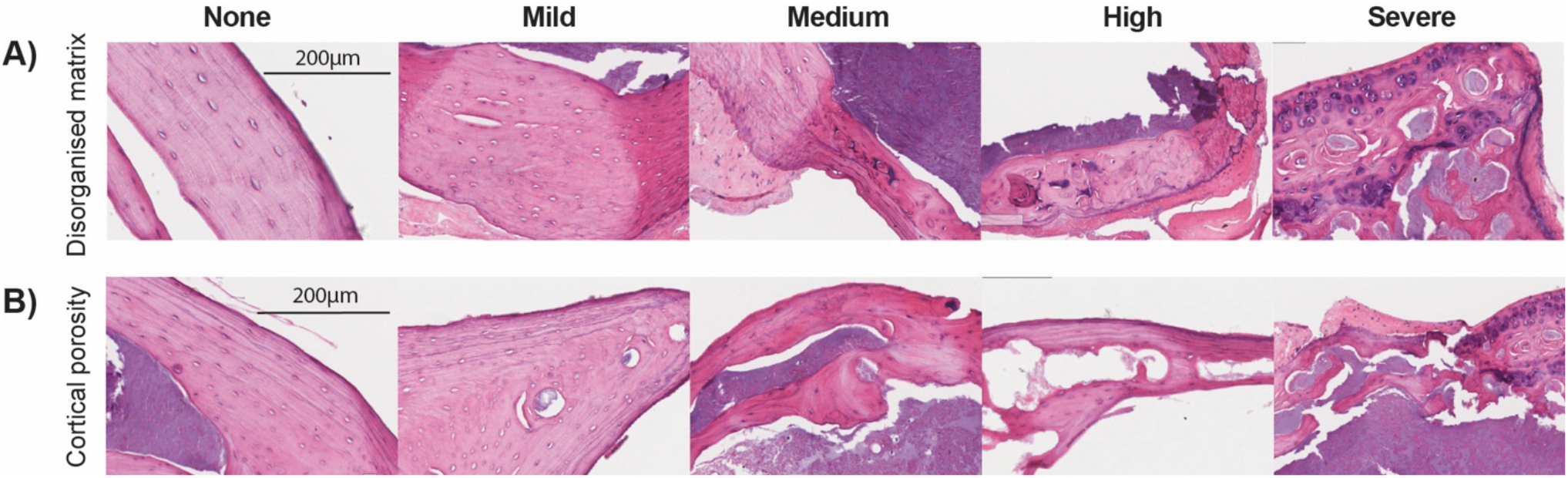
Histology scoring system. Example images for each category of scores for bone histology, for A) matrix disorganisation and B) cortical porosity.

### Ovarian analysis

Ovarian histology was assessed as described previously [5], and is described again below. To assess ovarian reserves, ovaries were collected from diestrus stage animals, embedded in resin, subject to thick (20 µm) sections for the entire ovary, with every third section digitally scanned for stereology analysis, and follicle reserve and health manually counted on a grid using Stereo Investigator software. Slides were labelled with a code that was separately assigned to each sample by an independent investigator, so that all analyses were conducted under blinded conditions. Detailed descriptions of each step of this analysis are provided below.

#### Estrous cycle tracking

Estrous cycles were tracked for five consecutive days to determine when mice were at diestrus, to allow for ovaries to be collected at a consistent stage of their cycle. Vaginal smears were performed using 20 µl of 0.9% saline and left to dry on slides. They were then stained with 0.5% toluidine blue, rinsed with water and examined under a light microscope. The presence and number of epithelial cells – cornified or nucleated, and leukocytes was assessed to determine the cycle stage, defined as: estrous – mostly cornified epithelial cells; metestrus – both cornified epithelial cells and leukocytes; diestrus – mostly leukocytes; proestrus – mainly nucleated, some cornified epithelial cells.

#### Tissue collection

When animals were confirmed to be at diestrus, they were subject to isofluorane euthanasia, and blood was collected by cardiac puncture using a 1 ml tuberculin syringe and 30-gauge needle (BD Medical, USA). Blood was then transferred to a micro centrifuge tube and left at room temperature to cloth for 30 minutes. Centrifugation was then performed at 5,000 *g* for 10 min. to collect serum, which was subsequently transferred to a new tube and stored at −80°C. After cardiac puncture, mice were immediately euthanized by cervical dislocation, and tissues collected. Ovaries were carefully cleaned from remaining fat under a dissecting microscope and weighed. One ovary from each animal was fixed in 4% paraformaldehyde (PFA) at 4°C overnight and transferred to 70% ethanol the next day for storage. The other ovary was placed in a micro centrifuge tube and snap frozen in liquid nitrogen for molecular analyses.

#### Resin embedding

Fixed ovaries stored in ethanol were dehydrated and embedded in resin with a kit by Technovit 7100 (Heraeus Kulzer, Wehrheim, Germany) according to the protocol provided by the manufacturer. The resin blocks containing the fixed ovarian samples were allowed to solidify and dry before they were taken out of the mould at three weeks. They were then sectioned at 20 µm on a Leica RM2252 microtome with a 16 cm d-profile TC knife. The cut sections were immersed in RO water, unfolded, straightened and placed on glass slides (Menzel-Glaser Superfrost Plus, Thermo Fisher Scientific, Sydney, NSW, Australia). All slides were placed on a heat plate at 37°C to dry and set and subsequently stored at room temperature.

#### Schiff staining of resin ovarian sections

The ovarian sections, previously mounted on slides and dried, were submerged in a bath of periodic acid (POCD Scientific, Artarmon, NSW, Australia) for 30 mins, rinsed with tap water for 5 mins, and stained with Schiff’s reagent (Fronine, Thermo Fisher Scientific, Soresby, VIC, Australia) for 45 mins at room temperature in a fume hood. Slides were rinsed with tap water for 5 mins and counterstained with filtered Mayer’s hematoxylin (Sigma-Aldrich, Castle Hill, NSW, Australia) for 2 hrs in an incubator at 37°C. Then slides were rinsed again with tap water for 5 mins, followed by Scott’s blue (POCD Scientific, Artarmon, NSW, Australia) stain in the fume hood for 3 mins at room temperature and washed in tap water for 3 mins. With this, staining was completed, and slides were left to dry overnight. They were put onto coverslips the next day using DPX mounting media for histology (Sigma Aldrich, Castle Hill, NSW, Australia) and coverslips (KNITTEL Coverglass, ProSciTech, Kirwan, Queensland, Australia). Each slide contained at least five serial sections, and was marked with a code that did not indicate treatments, allowing assessment to be performed in a blinded fashion.

#### Slide scanning

The Olympus VS200 Research Slide Scanner was used to acquire datasets for stereology analysis. Overview images of the whole slideweres captured with the PLAN 2x NA 0.06 objective. The UPLXAPO 40x NA 0.95 objective was used to acquire a detailed virtual z-stack with 1 μm increments of each section with 1 ms exposure. The captured 40x virtual z-stack was then loaded into MBF Stereo Investigator (MBF Bioscience, USA) for stereology analysis.

#### Stereology

Stereo Investigator (SI) software was used to obtain accurate ovarian follicle counts, as previously described [5, 74]. Scanned sections were imported as a virtual z-stack in SI with a cut thickness of 20 *μ*m and mounted thickness of 15 *μ*m. Every third consecutive section of each ovary was assessed by a blinded investigator. Lens calibration was performed to match the image resolution.

Large growing follicles, including secondary, small antral, large antral, and pre-ovulatory follicles were counted across the entire section. They were only included in the count if the nucleus was visible to prevent overcounting. Corpora lutea (CLs) were counted per ovary, while tracking across sections to confirm a CL. The results presented of those follicles is a raw count. By contrast, small follicles (primordial, transitory, and primary) were counted on a grid, randomly placed in the optical fractionator workflow. According to the workflow, the region of interest (all ovarian tissue) was selected under low magnification with the auto-trace assist. The grid size was then set to 800 and counting frame to 500. All follicles were counted within the square counting frame. The frame had two of its sides coloured red and two coloured green. For follicles lying on a line, only follicles lying on green lines were included in the count, while follicles crossed by the red line were excluded.

When all sections of a particular ovary were complete, they were put together to calculate the follicle population estimate for the entire ovary. This calculation accounts for area sampling fraction (counting frame/grid size), section sampling fraction (every third section), and mounted thickness over counting frame height, as per the software default. Unhealthy follicles were also assessed according to their size – large ones across the entire section and small ones on the grid.

All histology sample preparation, slide scanning, follicle counting and health assessments were performed in a blinded fashion, with samples labelled with codes that were unknown to the investigator analysing each sample. Data were recorded with codes only, and unblinding to experimental groups for each sample only occurred once all data had been recorded.

#### Follicle classification

Morphological classification of the ovarian follicles was performed according to previously defined standards [75]. Primordial follicles were described by an oocyte surrounded by a single layer of squamous granulosa cells (GCs). The transition stage between primordial and primary follicle is defined by an oocyte and a single layer of GCs, some of which can be squamous and some cuboidal. Primary follicles consist of an oocyte and a single layer of cuboidal GCs. Primordial follicles are described as “non-growing” or the “ovarian reserve”, and transitory and primary as “small growing” follicles. These can be presented as a ratio of growing/non-growing to illustrate follicle activation rate [76]. This ratio was calculated by summing the transitory and primary follicle estimations together and dividing them by the primordial follicle estimation.

Secondary follicles are described as having a growing oocyte and at least two complete layers of cuboidal GCs. Antral follicles are defined by two or more GC layers and an antral space. Depending on the volume of the antral space, the antral follicle can be classified as small antral or large antral. A pre-ovulatory follicle is a large antral right before ovulation and is characterised by the oocyte and cumulus cells around it being surrounded by antral space from all sides, apart from a thin stalk of cumulus cells adhering it to the follicle. These large growing follicles were enumerated independently and in combination: “all antral” being small, large antral and pre-ovulatory; and “all big follicles” including all antral and secondary follicles.

Unhealthy follicles were also quantified. They were classified according to defined morphologies and put into categories: atretic (>10% pycnotic GCs or damaged oocyte), multinuclear, vacuoles present, enlarged oocyte with undifferentiated GCs, biovular follicles, and zona pellucida remnants (ZPRs). With the exception of ZPRs, these were grouped as “large unhealthy follicles” and were counted across the whole section. The level of follicle/oocyte damage was expressed as a healthy/unhealthy follicle ratio. Health ratios (proportions) were calculated as healthy/total for each subgroup (primordial, transitory, and primary), with max value of 1.

Small unhealthy follicles were counted on the grid (as described above), in order to be comparable to the healthy small follicles. They were defined by missing oocyte or ZPRs still surrounded by a single layer of GCs. Depending on the GCs (squamous or cuboidal) they were described as unhealthy primordial, transitory or primary. This was used for a specific healthy/unhealthy ratio for each small follicle type.

### Steroid hormone measurements

Steroid hormone profiles for estradiol (E2), estrone (E1), progesterone (P4), testosterone (T), and androstenedione (A4) were quantified from serum using a ultrasensitive LC-MSMS assay [77] following the derivatization of estradiol [78], and performed as contract research in the lab of Prof David Handelsmann (ANZAC medical research institute, Concord, NSW Australia). Briefly, serum samples collected in a volume of 0.2 ml were subject to LC-MSMS and calibrated against certified reference steroid hormone materials.

### Serum analytes

Inorganic phosphate was measured using a malachite colourimetric assay from Sigma (MAK030). Parathyroid hormone (PTH) was measured using a two-site ELISA against mouse PTH 1-84 from Immutopics (now QuidelOrtho), catalogue number P160-2305. The MILLIPLEX Mouse Pituitary Magnetic Bead Panel (MPTMAG-49K) was used to measure follicle stimulating hormone, growth hormone, prolactin, thyroid stimulating hormone and lutenising hormone. Blood urea nitrogen was measured using a kit from Invitrogen (now ThermoFisher Scientific, catalogue number EIABUN). ELISA kits were used to measure cystatin C (Invitrogen EMCST3), Klotho (Cusabio CSB-E14362m) and FGF23 (Abcam ab213863). Elemental phosphorus and calcium were measured by ICP elemental analysis following the digestion of serum samples with concentrated nitric acid and heating.

### Behaviour

The elevated plus maze (EPM) was carried out during the light phase over 10 minutes, using a plus shaped maze consisting of 2 opposing open arms (50×10 cm) and 2 closed arms (50×10 cm). Behaviour was analysed by video and recorded as number of entries into the open and closed arms, time spent in the arms and time spent immobile. Data are presented as time spent in the open arms (%), open arm entries (% total entries), distance travelled (m) and time spent immobile (s).

### Statistics

Experiments were planned as 2×2 factorial designs, with data subject to mixed linear model analysis in *R* (version 4.2.1) and tests for the impact of NMN treatment within chemotherapy or vehicle control calculated by estimated marginal means using base *R* and the package *emmeans* (version 1.8.0), with p-values then subject to a Bonferroni correction for multiple comparisons. Data are summarised in figures as Tukey boxplots, showing the 25-75% interquartile range with whiskers indicating 95% confidence intervals, mean values indicated by a line within boxplots. In Fig 3, bone matrix disorganisation and cortical porosity were assessed by creating an ordinal logistic regression model, followed by estimated marginal means and Bonferroni-corrected t-tests. Detailed, annotated *R* scripts and raw CSV files used to generate statistical analyses and generate figures have been uploaded as supplementary files, allowing reproducible analysis – please see data availability section.

### Study design

This study was conducted in accordance with the ARRIVE guidelines on the use of animals in research.

The experimental unit for all experiments were data from individual animals, including parameters for each bone for each animal (Fig. 1-3, 5), ovarian reserve per ovary (Fig. 5), plasma analytes (Figs. 4, 6) and behaviour (Fig. 7).

The inclusion criteria for the study were female animals treated with CDDP and NMN as described above. Exclusion criteria were if animals had to be euthanased from the study due to poor health – a key requirement of our ethics approval – prior to reaching old age. Notably, studies of aged mice (Figs. 1-3) started with n=10 per group, however some groups had lower numbers than this due to animals being euthanased prior to reaching old age. This was particularly the case for animals treated with CDDP alone. Further, some bone samples were excluded for analysis due to damage occurring during tissue cleaning and preparation, with the initial µ-CT scan showing additional cases where bones had already been broken, and were therefore not subject to mechanical testing (Fig. 2) – further details above.

Animals were randomised into chemotherapy or vehicle treatment based on the order that they were first picked up out of the cage, alternating between the two for the next animal. Cages were similarly randomised into NMN or vehicle treatment in their drinking water based on the order they were taken from the rack, alternating between each intervention. This strategy of alternating treatments between each subsequent animal or cage was used to minimise the likelihood of confounding effects.

As described in previous sections, investigators were blinded to treatment groups at the time of tissue collection and during histological assessment of ovarian reserve, bone structure, and during mechanical testing and µ-CT analysis. Cages were labelled with animal numbers, and treatment groups correlating to animal numbers were recorded in a separate lab book.

Data were analysed by mixed linear model analysis or ordinal logistic regression modelling followed by posthoc analysis of estimated marginal means, as described in the statistics section above. We did not adjust this model for data that did not meet tests for normality, as mixed linear models are generally robust to violations of this assumption [79].

Sample size was determined based on precedents from previous papers in the field that have used similar primary outcomes in models of chemotherapy induced infertility [21, 22, 80, 81].

The primary outcome of each experiment is described in the relevant figure legend, with additional detail in the methods section above.

### Figures

Data figures were generated in R using the packages *ggplot2* (3.4.0), *ggtext* (0.1.2), *ggpubr* (0.4.0), *ggsignif* (0.6.3) and *ggh4x* (0.2.3.9000), with a custom script based on *rstatix* for the automatic placement of significance values. Data are shown as Tukey boxplots, with all raw data points overlaid. The illustration in Figure 1 was generated under license using BioRender.com.

## Data availability

All raw data have been uploaded to our Mendeley data site (reserved DOI: 10.17632/d684g7xzk6.1) as .csv files, with annotated R scripts for processing data, statistical analysis and generating figures. For µ-CT scans and digital scans of histological assessment of ovarian reserve, these very large files have not been uploaded due to file size limits on Mendeley Data. These have been maintained for long-term storage on UNSW servers and are available upon request.

## ACKNOWLEDGEMENTS

The salary of LEW is supported from a Hevolution / American Federation for Aging Research (AFAR) New Investigator award. This work was supported by the National Health and Medical Research Council (NHMRC) of Australia, through grants APP1103689 and APP1122484 to LEW, DAS and HAH, APP1139763 to RBG, LEW and KAW, a Career Development Fellowship to LEW (APP1127821) and Senior Research and Investigator Fellowships to RBG (APP1117538 and APP2009940). We gratefully acknowledge assistance from the UNSW Biological Resource Centre and Tzong-Tyng Hung from the Biological Resource Imaging Laboratory (BRIL) of the Mark Wainwright Analytical Centre, UNSW, which is a member of the National Imaging Facility, and Rabeya Akter of the Inductively Coupled Plasma Lab of the Mark Wainwright Analytical Centre at UNSW. We also wish to thank the Solina Chau foundation and anonymous donors for their philanthropic support.

